# GLP-1R agonist NLY01 reduces retinal inflammation, astrocyte reactivity, and retinal ganglion cell death secondary to ocular hypertension

**DOI:** 10.1101/2020.06.18.146720

**Authors:** Jacob K. Sterling, Modupe Adetunji, Samyuktha Guttha, Albert Bargoud, Katherine Uyhazi, Ahmara G. Ross, Joshua L. Dunaief, Qi N. Cui

**Author notes:** Correspondence should be addressed to Qi N. Cui, M.D., Ph.D.

## Abstract

Glaucoma is the leading cause of irreversible blindness worldwide and is characterized by the death of retinal ganglion cells. Reduction of intraocular pressure (IOP) is the only therapeutic mechanism available to slow disease progression. However, glaucoma can continue to progress despite normalization of IOP. New treatments are needed to reduce vision loss and improve outcomes for patients who have exhausted existing therapeutic avenues. Recent studies have implicated neuroinflammation in the pathogenesis of neurodegenerative diseases of both the retina and the brain, including glaucoma and Parkinson’s disease. Pro-inflammatory A1 astrocytes contribute to neuronal cell death in multiple disease processes and have been targeted therapeutically in mouse models of Parkinson’s disease. Microglial release of pro-inflammatory cytokines C1q, IL-1α, and TNF-α is sufficient to drive the formation of A1 astrocytes. The role of A1 astrocytes in glaucoma pathogenesis has not been explored. Using a mouse model of glaucoma, we demonstrated that IOP elevation was sufficient to trigger production of C1q, IL-1α, and TNF-α by infiltrating macrophages followed by resident microglia. These three cytokines drove the formation of A1 astrocytes in the retina. Furthermore, cytokine production and A1 astrocyte transformation persisted following IOP normalization. Ablation of this pathway, by either genetic deletions of C1q, IL-1α, and TNF-α, or treatment with glucagon-like peptide-1 receptor agonist NLY01, reduced A1 astrocyte transformation and RGC death. Together, these results highlight a new neuroinflammatory mechanism behind glaucomatous neurodegeneration that can be therapeutically targeted by NLY01 administration.

## INTRODUCTION

Glaucoma is characterized by the death of retinal ganglion cells (RGCs) leading to permanent vision loss. It is the leading cause of irreversible blindness globally and is projected to affect approximately 112 million people worldwide by 2040 (Tham et al., 2014). Elevated intraocular pressure (IOP) is strongly associated with glaucoma, and reduction of IOP is the only therapeutic mechanism available to slow disease progression. However, glaucoma can continue to progress even in patients who achieve normal IOPs following medical and/or surgical treatments (Quigley, 2018). New therapies are urgently needed to prevent vision loss in patients suffering from glaucoma.

Reactive astrocytes are observed in multiple neurodegenerative diseases (Liddelow et al., 2017; Yun et al., 2018). In healthy neural tissue, astrocytes serve a wide variety of roles. They contribute to neurotransmitter recycling, neuronal metabolism, and formation of the blood-brain and blood-retina barriers (Clarke and Barres, 2013; Liddelow and Barres, 2017). In the retina, astrocytes are found exclusively in the ganglion cell layer, comingled with RGCs (Vecino et al., 2016). In response to both local and systemic stimuli, astrocytes can adopt reactive forms, A1 pro-inflammatory or A2 neuroprotective, both of which have been transcriptionally defined. A1 reactive astrocytes lose their phagocytic capacity as well as their ability to promote synapse formation and function. At the same time, A1 astrocytes gain pro-inflammatory and neurotoxic functions (Liddelow et al., 2017; Zamanian et al., 2012). In contrast, A2 astrocytes, observed in post-ischemic tissue, upregulate neurotrophic factors, promoting a neuroprotective environment (Zamanian et al., 2012). While A1 astrocytes have been implicated in multiple neurodegenerative diseases (Liddelow et al., 2017; Yun et al., 2018), the reactive states of astrocytes and their contribution in glaucoma is not known.

In the brain and the retina, neurotoxic A1 astrocytes are induced by microglial release of pro-inflammatory cytokines IL-1α, TNF-α, and C1q (Liddelow et al., 2017). Strong links exist between these three cytokines and glaucoma. *IL1A* and *TNF* polymorphisms are associated with primary open angle glaucoma (Bozkurt et al., 2011; Fan et al., 2010; Mookherjee et al., 2010; Wang et al., 2006). TNF-α protein levels are elevated in the vitreous, retina, and optic nerves of glaucomatous eyes (Williams et al., 2017). In the DBA/2J mouse model of hypertensive glaucoma, *C1qa* mRNA levels are associated with disease progression (Stevens et al., 2007) and C1q inhibition is sufficient to prevent early RGC synapse loss and RGC death (Howell et al., 2011, 2014; Williams et al., 2016). Multiple publications have also demonstrated C1q upregulation in glaucomatous human eyes (Reinehr et al., 2016; Stasi et al., 2006). Although IL-1α, TNF-α and C1q have been independently implicated in glaucoma, it is not known whether they induce A1 astrocyte reactivity in glaucomatous retinas.

Glucagon-like peptide 1 (GLP-1) is an incretin hormone that regulates blood glucose, weight, and satiety through its action at the GLP-1 receptor (GLP-1R) in both the systemic circulation and the central nervous system (Drucker, 2018). NLY01 is a long-acting GLP-1R agonist with an extended half-life and favorable blood-brain barrier penetration (Yun et al., 2018). In mouse models of Parkinson’s disease (PD), A1 astrocytes contribute to dopaminergic cell death and poor motor phenotypes. NLY01 has been shown to reduce microglial production of C1q, TNF-α, and IL-1α, thereby blocking A1 astrocyte transformation, reducing dopaminergic cell death, and improving motor symptoms in mouse models of PD (Yun et al., 2018).

Using the microbead-induced ocular hypertension mouse model of glaucoma, we showed that microglia and infiltrating macrophages upregulated C1q, TNF-α, and IL-1α. These three cytokines were necessary for A1 transformation within the retina. Cytokine upregulation and A1 transformation persisted even after IOP returned to normal levels. Genetic deletion of these cytokines prevented A1 astrocyte formation and RGC loss. Finally, NLY01 therapy reduced microglial and macrophage activation, A1 astrocyte transformation, and RGC loss in our model. Together, these data demonstrate that GLP-1R activation is capable of reducing ocular inflammation driven by both resident microglia and infiltrating macrophages, thereby preventing A1 astrocyte activation and rescuing RGCs in ocular hypertension glaucoma. NLY01 has potential for clinical use in the treatment of glaucoma and possibly other retinal diseases characterized by deleterious glial reactivity.

## RESULTS

### Elevated intraocular pressure induces A1 astrocyte reactivity in the retina

Magnetic microbeads (left eye) or balanced saline solution (BSS, right eye) were injected into the anterior chamber (AC) of wild-type (WT) and *Il1a−/−*; *Tnf−/−*; *C1qa−/−* triple knockout (TKO) mice. Intraocular pressure (IOP) was recorded at 1, 2, 3, 5, and 7 days post-injection, and weekly thereafter. Microbead-injected eyes (Bead) in both WT and TKO animals had elevated intraocular pressures (eIOP) beginning at 1 week post-injection compared to BSS-injected eyes (BSS) (Fig. 1A). IOPs peaked at 2 weeks post-injection and remained elevated through 35 days post-injection. There was no difference in IOP between Bead and BSS eyes by 42 days post-injection (Fig. 1A). Neurosensory retinas were isolated for cell sorting at 3, 14, and 42 days post-injection (p.i.) (Fig. S1A). Cell-type enrichment for all fractions used in the paper were validated by qPCR (Fig. S1B-C). Astrocytes and Müller cells were isolated using ASCA2+ selection, as reported previously (Kantzer et al., 2017). By 14 days post-injection, ACSA2+ cells exhibited increased levels of pan reactive and A1 astrocyte specific markers (Fig. 1B-D), suggesting that A1 astrocyte transformation occurs early in the disease process. A1 reactivity persisted in microbead-injected WT animals at 42 days post-injection, despite normalization of IOP (Fig. 1D). TKO animals failed to form A1 astrocytes (Fig. 1C-D) despite eIOP (Fig. 1A). This was consistent with previous findings (Liddelow et al., 2017). Markers of A2 reactivity were consistently unchanged in WT microbead-injected eyes compared to WT BSS-injected eyes (Fig. 1C-D).

**Figure 1.**
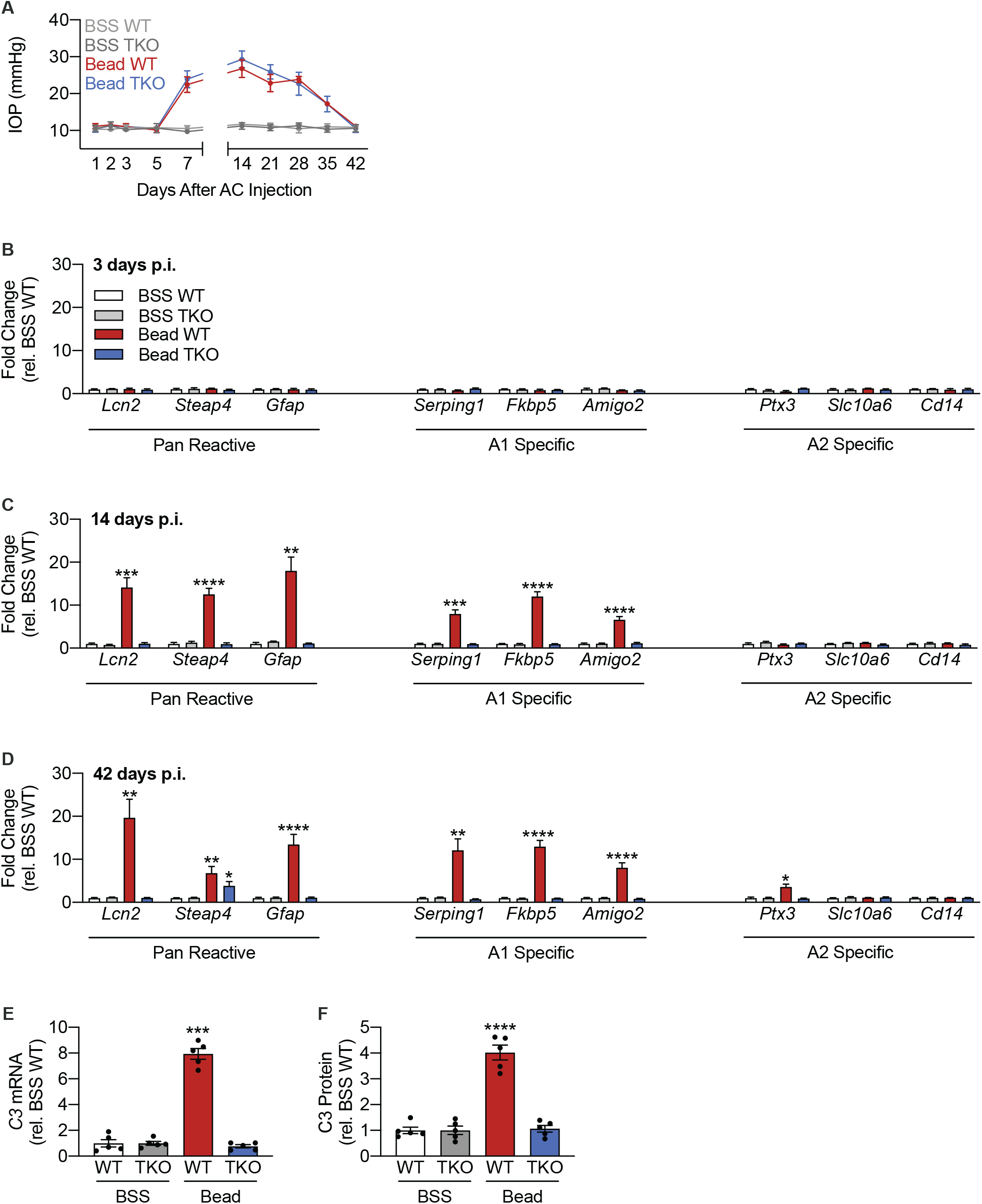
Elevated intraocular pressure induces A1 astrocyte reactivity in the retina. C57BL6/J (WT) and *Il1a−/−*; *Tnf−/−*; *C1qa−/−* knockout (TKO) mice were injected either with microbeads (“Bead”, left eye) into the anterior chamber (“AC”), to increase intraocular pressure (IOP), or with BSS (right eye). (A) IOP measurements across the duration of the study in both eyes of WT and TKO animals. (n=20 eyes per condition per genotype) (B-D) qPCR measurements of pan reactive, A1-specific, and A2 specific transcripts from ACSA2+ cells isolated from WT and TKO mice at 3 days (B), 14 days (C), and 42 days (D) post-injection (“p.i.”). (n=3 eyes per condition per genotype) (E) qPCR measurements of *C3* mRNA levels in ACSA2+ cells. (n=5 eyes per condition per genotype) (F) ELISA measurements of C3 protein levels in ACSA2+ cells. (n=5 eyes per condition per genotype) All data presented as mean ± SEM. Mann-Whitney U test, *p<0.05, **p<0.01, ***p<0.001, ****p<0.0001 vs. BSS WT. See also Figure S1.

Liddelow *et al.* demonstrated that complement component 3 (C3) is a marker for A1 astrocytes (Liddelow et al., 2017). We measured C3 mRNA and protein levels in ACSA2+ cells isolated from both BSS- and microbead-injected WT and TKO retinas. C3 mRNA (Fig. 1E) and protein (Fig. 1F) levels were elevated in WT microbead-injected eyes compared to both BSS-injected eyes and TKO microbead-injected eyes (Fig. 1E-F). These data suggest that C3 production in ACSA2+ cells, which encompass both astrocytes and Müller cells, is dependent on IL-1α, TNF-α, and C1q, and that loss of A1 reactivity reduces C3 production.

### A1 astrocytes contribute to retinal ganglion cell death secondary to eIOP

To determine whether A1 astrocytes play a role in retinal ganglion cell (RGC) death in the microbead-induced eIOP model of glaucoma, WT, *Il1a−/−*; *Tnf−/−* double knockout (DKO) mice, *C1qa−/−* single knockout mice, and TKO mice were injected with magnetic microbeads in one eye and BSS in the fellow eye. After 42 days, whole retina flatmounts were stained for the RGC marker Brn3a (Nadal-Nicolás et al., 2009). There was no difference in RGC loss observed in *Il1a−/−*; *Tnf−/−* DKO mice compared to WT mice. There was a modest improvement in RGC survival observed in *C1qa−/−* mice compared to WT mice, consistent with previously published results (Howell et al., 2011, 2014; Williams et al., 2016). RGC death was reduced in TKO mice compared to all other genotypes (Fig. 2A), suggesting that the loss of all three cytokines provided an additional benefit beyond the loss of either IL-1α and TNF-α or C1q alone. Together these data suggest that A1 astrocytes contribute to RGC death caused by eIOP.

**Figure 2.**
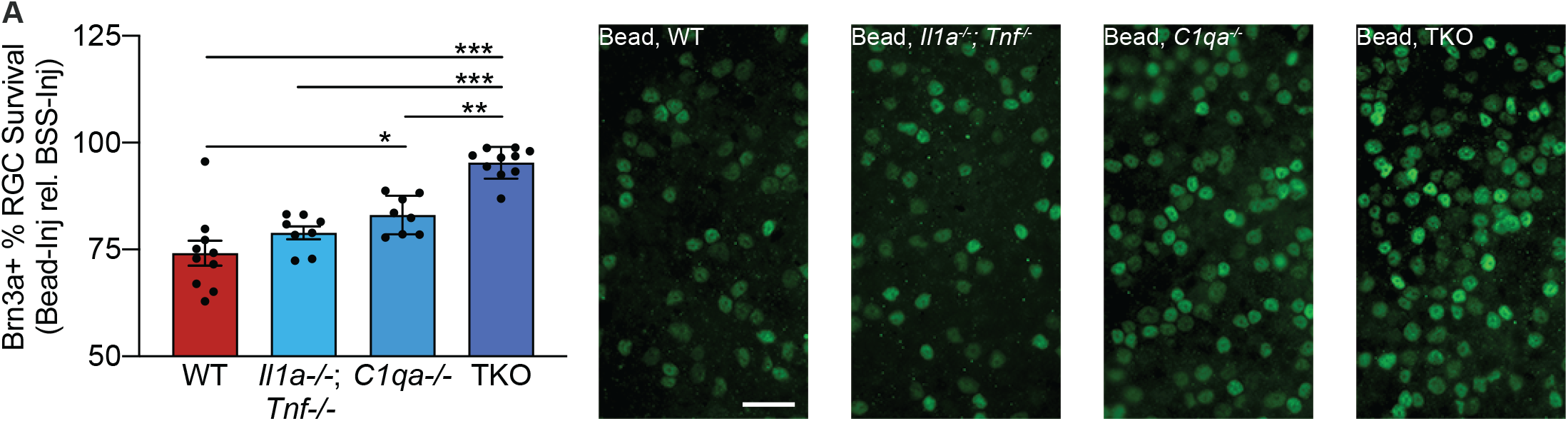
A1 astrocytes trigger retinal ganglion cell death secondary to eIOP. (A) C57BL6/J (WT), *Il1a−/−*; *Tnf−/−*, *C1qa−/−* and *Il1a−/−*; *Tnf−/−*; *C1qa−/−* knockout (TKO) mice were injected either with microbeads (left eye) into the anterior chamber (“AC”), to increase intraocular pressure (IOP), or with BSS (right eye). At 42 days post-injection mice were euthanized and retinal flatmounts were stained for the retinal ganglion cell (RGC) marker Brn3a. Cell count for each microbead-injected eye was normalized against the contralateral BSS-injected eye to determine percent survival of Brn3a+ cells. n=10 mice per genotype. All data presented as mean ± SEM. One way ANOVA with Tukey’s multiple comparisons test, *p<0.05, **p<0.01, ***p<0.001. Representative images are shown on the right. Scale bar = 50μm.

### Early retinal inflammation is driven by CD11b+ CD11c+ cells and persists beyond re-normalization of IOP

Although IL-1α, TNF-α, and C1q are necessary for A1 astrocyte transformation secondary to eIOP (Fig. 1), the time course and source of IL-1α, TNF-α, and C1q production in this model were not known. To address both of these questions, we injected either microbeads or BSS into the anterior chamber of WT mice. Neurosensory retinas were isolated at 1, 2, 3, 7, 14, 28 and 42 days after injection and used either for ELISA to measure IL-1α, TNF-α, and C1q protein levels, or dissociated for cell sorting. IL-1α, TNF-α, and C1q protein levels increased in line with IOP, rising by day 7 and then plateauing at subsequent time points (Fig. 3A-C). IL-1α, TNF-α, and C1q remained elevated at 6 weeks post-injection, despite a return of IOP to baseline levels (Fig. 1A, Fig. 3A-C). Microbead-injected eyes that did not have a significant increase in IOP, observed in approximately 5% of eyes, did not have elevated IL-1α, TNF-α, or C1q mRNA levels in CD11b+ cells 42 days after injection (Fig. S2A-C). Together these data suggest that the retinal inflammatory response, once initiated, outlasted IOP elevation.

**Figure 3.**
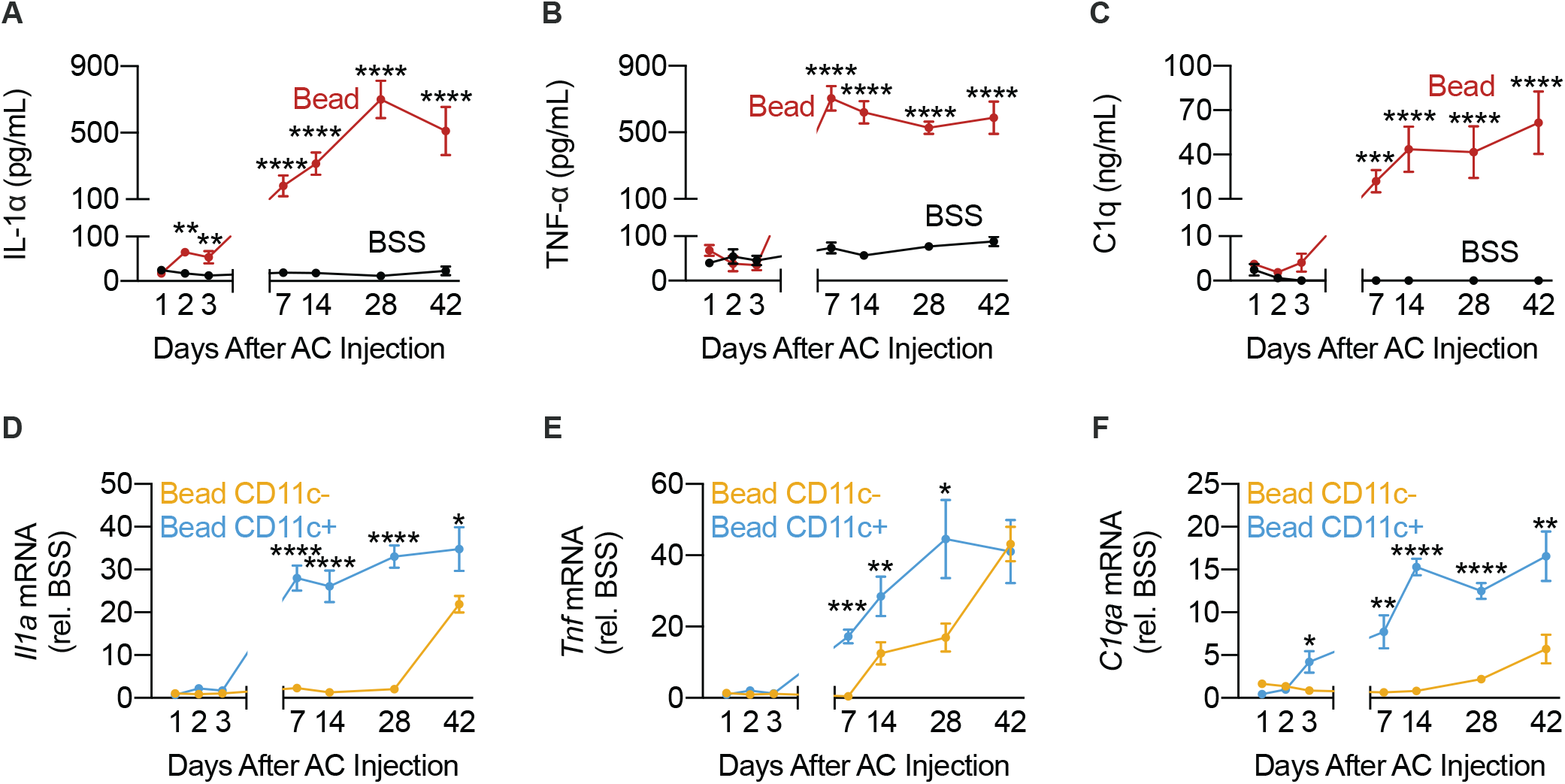
Early retinal inflammation is driven by CD11b+ CD11c+ cells and outlasts re-normalization of IOP. C57BL6/J (WT) mice were injected either with microbeads (“Bead”, left eye) into the anterior chamber (“AC”), to increase intraocular pressure (IOP), or with BSS (right eye). (A-C) ELISA measurements of IL-1α (A), TNF-α (B), or C1q (C) protein levels in whole neurosensory retina at 1, 2, 3, 5, 7, 14, 28, and 42 days post-injection. Statistical test compared BSS-injected eyes to microbead-injected eyes at the same time points. (n=5 eyes per time point per condition) (D-F) CD11b+ CD11c− and CD11b+ CD11c+ cells were isolated from neurosensory retina at 1, 2, 3, 5, 7, 14, 28, and 42 days post-injection. *Il1a* (D), *Tnf* (E), and *C1qa* (F) mRNA levels were measured by qPCR in both cell populations. qPCR measurements in microbead-injected eyes were normalized to the contralateral BSS-injected eyes. Statistical tests compared microbead-injected CD11b+ CD11c+ cells and microbead-injected CD11b+ CD11c− cells at the same time points. (n=5 eyes per time point per condition) All data presented as mean ± SEM. Mann-Whitney U test, *p<0.05, **p<0.01, ***p<0.001, ****p<0.0001. See also Figure S2.

In other mouse models of neuroinflammation, resident microglia are responsible for the production of IL-1α, TNF-α and C1q, leading to A1 astrocyte transformation (Liddelow et al., 2017; Yun et al., 2018). However, recent work in the DBA/2J mouse model of glaucoma demonstrated that resident retinal microglia (CD11b+ CD11c−) adopted an anti-inflammatory state early on in their response to eIOP. The authors suggested that CD11b+ CD11c+ cells, a population enriched with infiltrating macrophages, contributed to early eIOP-induced inflammation (Tribble et al., 2019). We compared the expression of *Il1a*, *Tnf*, and *C1qa* between CD11b+ CD11c− (predominantly microglia) and CD11b+ CD11c+ (predominantly macrophages) cells isolated from microbead-injected eyes. Both sets of microbead-injected eyes were normalized against BSS-injected eyes. CD11b+ CD11c+ cells exhibited earlier expression of *Il1a*, *Tnf*, and *C1qa* compared to CD11b+ CD11c− cells (Fig. 3D-F). Expression of *Il1a*, *Tnf*, and *C1qa* by CD11b+ CD11c− cells also increased but lagged behind those of CD11b+ CD11c+ cells by days to weeks (Fig. 3D-F). These results suggest that infiltrating macrophages provided the primary source of pro-inflammatory signals in early glaucomatous retinal inflammation following eIOP. In further support of this hypothesis, we demonstrated that A1 astrocytes were present by 14 days post-injection (Fig. 1B-D), when CD11b+ CD11c− cells did not express all three cytokines necessary for A1 transformation (Fig. 3D-F). Therefore, A1 transformation could not have been driven by CD11b+ CD11c− cells in this model.

### NLY01, a GLP-1R agonist, reduces IL-1α, TNF-α, and C1q production by CD11b+ CD11c+ and CD11b+ CD11c− cells during eIOP, thereby reducing A1 astrocyte activation

NLY01, a glucagon-like peptide 1 receptor (GLP-1R) agonist, has previously been shown to modulate microglial phenotype resulting in reduced A1 astrocyte activation in the brain (Yun et al., 2018). We hypothesized that NLY01 therapy would reduce IL-1α, TNF-α, and C1q production by both microglia and macrophages, and thereby decrease A1 astrocyte conversion secondary to eIOP. To test the efficacy of NLY01 in our model, we used microbead injections to induce eIOP. Mice were given twice weekly subcutaneous injections of either NLY01 at a dose of 5mg/kg or normal saline solution (NSS). Neurosensory retinas were harvested at 14 and 42 days after injection to evaluate the effects of NLY01 on both the CD11b+ CD11c+ mediated early response and the CD11b+ CD11c− mediated late response. NLY01 had no effect on intraocular pressure in microbead- or BSS-injected eyes (Fig. S3). By 14 days post-injection, NLY01 therapy reduced CD11b+ CD11c− (resident microglia enriched) production of Tnf-α (Fig. 4B) without altering basal expressions of IL-1α or C1q (Fig. 4A, 4C). NLY01 also reduced CD11b+ CD11c+ (infiltrating macrophage enriched) expression of IL-1α, TNF-α, and C1q (Fig. 4D-F). In the ACSA2+ fraction (astrocyte and Müller cell enriched), NLY01 also reduced expression of pan reactive transcripts, A1-specific transcripts, and C3, consistent with decreased A1 activation (Fig. 4G-H). By 42 days post-injection, NLY01 therapy reduced both CD11b+ CD11c− (resident microglia enriched) and CD11b+ CD11c− (infiltrating macrophage enriched) expression of IL-1α, TNF-α, and C1q (Fig. 5A-F). NLY01 also reduced expressions of pan reactive transcripts, A1-specific transcripts, and C3 in the ACSA2+ fraction (Fig. 5G-H), as it had by the 14-day post-injection time point (Fig. 4G-H).

**Figure 4.**
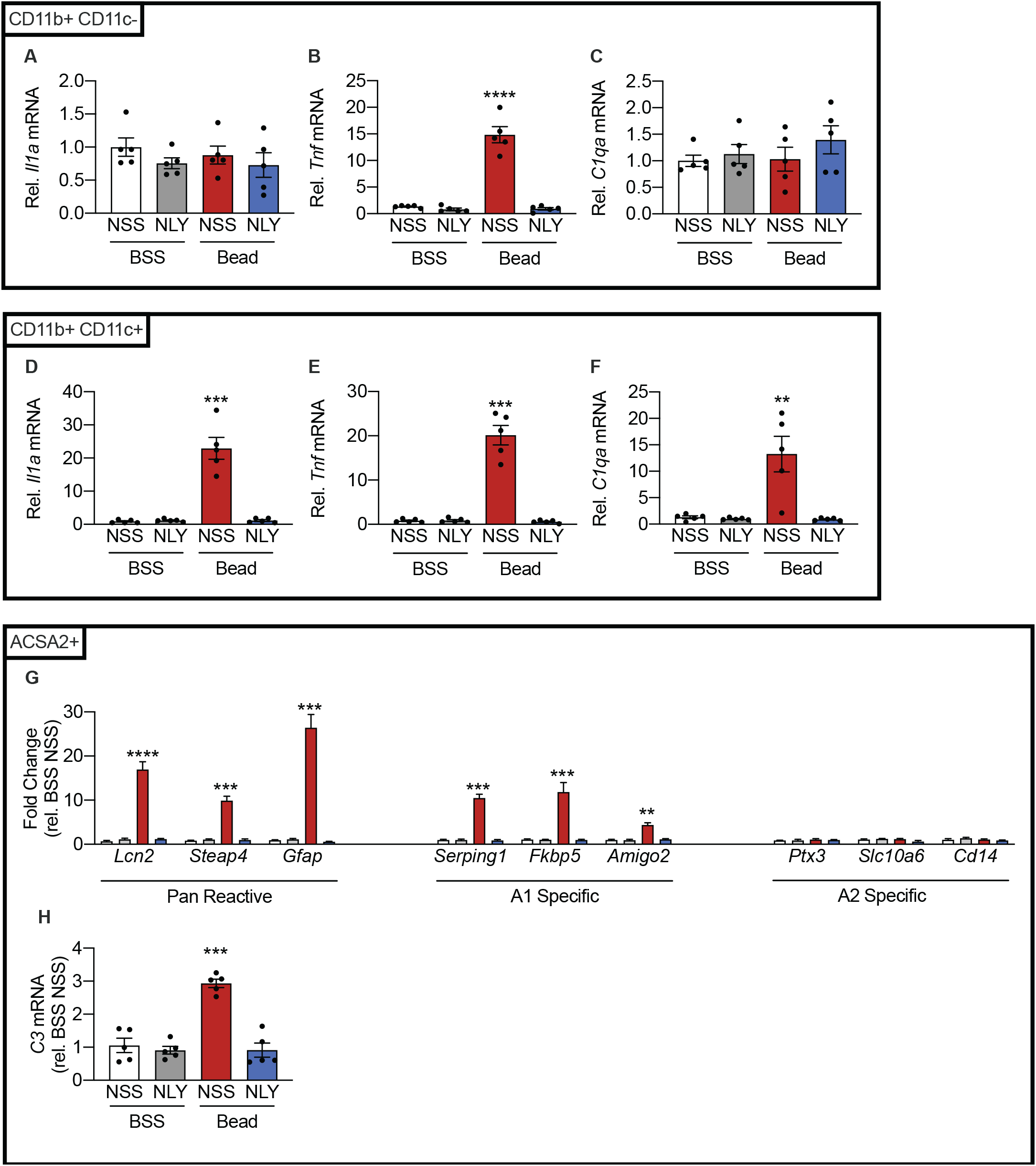
GLP-1R agonist, NLY01, reduces acute production of IL-1α, TNF-α and C1q secondary to eIOP and A1 astrocyte activation. C57BL6/J (WT) mice were injected either with microbeads (“Bead”, left eye), to increase intraocular pressure (IOP), or with BSS (right eye). Following intraocular injections, mice were randomized to twice weekly subcutaneous NLY01 (5 mg kg^−1^ per injection) or normal saline. Mice were euthanized 14 days post-injection. (A-F) CD11b+ CD11c− and CD11b+ CD11c+ cells were isolated from neurosensory retina. qPCR was performed to measure *Il1a* (A, D), *Tnf* (B, E) and *C1qa* (C,F) mRNA levels in each population. (n=5 eyes per condition) (G) qPCR measurements of pan reactive, A1-specific and A2 specific transcripts from ACSA2+ cells 14 days post-injection. (n=5 eyes per condition) (H) qPCR measurement of *C3* mRNA levels in ACSA2+ cells. (n=5 eyes per condition) All data presented as mean ± SEM. Mann-Whitney U test, *p<0.05, **p<0.01, ***p<0.001, ****p<0.0001 relative to BSS NSS. See also Figure S3.

**Figure 5.**
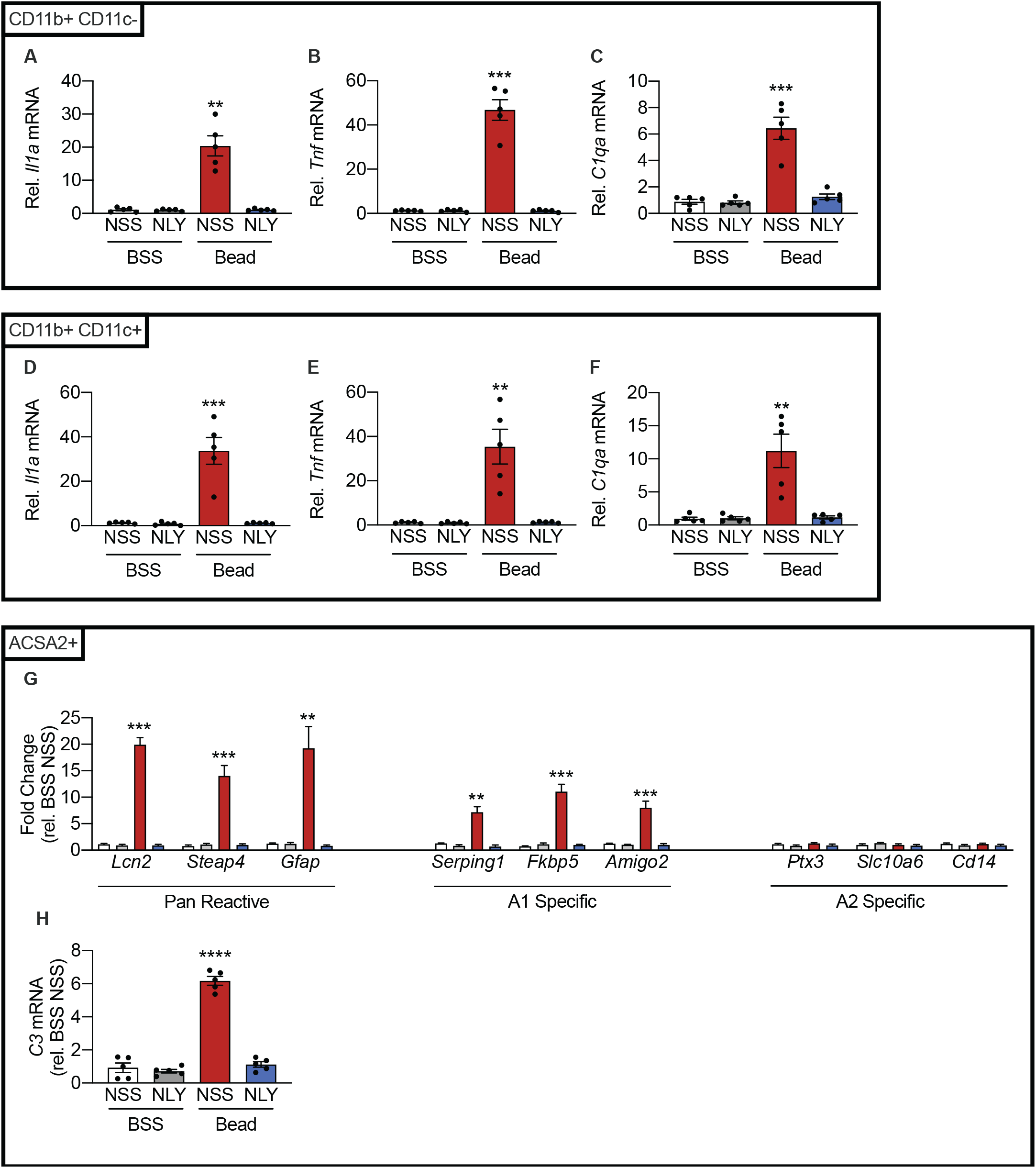
NLY01 reduces CD11b+ CD11c+ and CD11b+ CD11c− IL-1α, TNF-α and C1q production due to prolonged eIOP, reducing A1 astrocyte activation. C57BL6/J (WT) mice were injected either with microbeads (“Bead”, left eye), to increase intraocular pressure (IOP), or with BSS (right eye). Following injections, mice were randomized to twice weekly subcutaneous NLY01 (5 mg kg^−1^ per injection) or normal saline. Mice were euthanized 42 days post-injection. (A-F) CD11b+ CD11c− and CD11b+ CD11c+ cells were isolated from neurosensory retina. qPCR was performed to measure *Il1a* (A, D), *Tnf* (B, E), and *C1qa* (C,F) mRNA levels in each population. (n=5 eyes per condition) (G) qPCR measurements of pan reactive, A1-specific, and A2 specific transcripts from ACSA2+ cells 42 days post-injection. (n=5 eyes per condition) (H) qPCR measurement of *C3* mRNA levels in ACSA2+ cells. (n=5 eyes per condition) All data presented as mean ± SEM. Mann-Whitney U test, **p<0.01, ***p<0.001, ****p<0.0001 vs. BSS NSS. See also Figure S3.

### NLY01 reduces RGC death secondary to eIOP

To test the efficacy of NLY01 as a potential neuroprotective agent in our model, eIOP was once again induced through microbead injections in mice treated with either twice weekly NLY01 at a dose of 5 mg/kg or NSS. After 42 days, neurosensory retinas were isolated, flat-mounted, and labeled with RGC markers Brn3a (Fig. 6A) and Rbpms (Fig. 6B) for RGC counting. NLY01 therapy reduced RGC death secondary to eIOP as quantified by both labeling methods (Fig. 6A-B).

**Figure 6.**
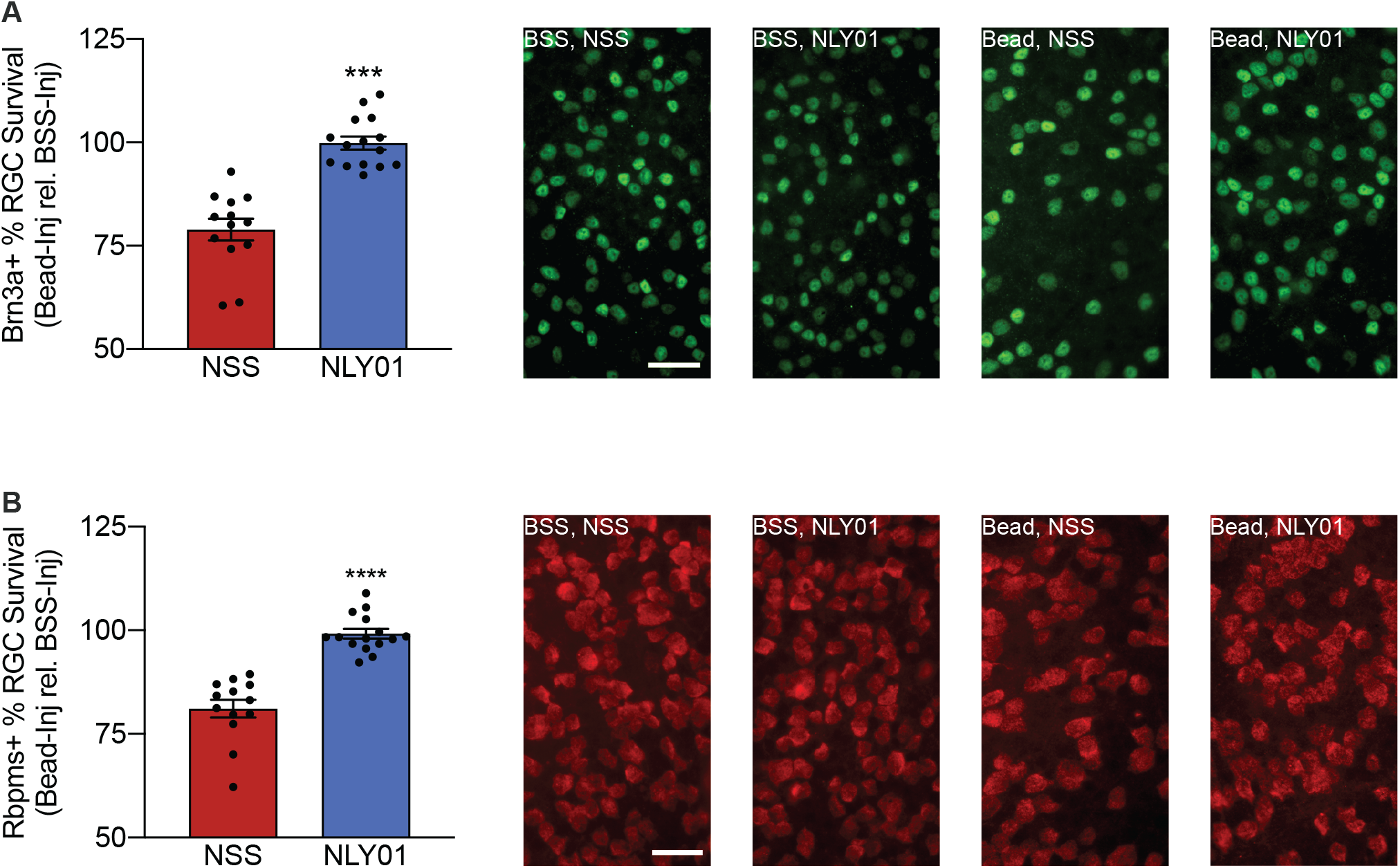
NLY01 reduces RGC death secondary to eIOP. C57BL6/J (WT) mice were injected either with microbeads (“Bead”, left eye), to increase intraocular pressure (IOP), or with BSS (right eye). Following injection, mice were randomized to twice weekly subcutaneous NLY01 (5 mg kg^−1^ per injection) or normal saline. Mice were euthanized 42 days post-injection. (A) Retinal flatmounts were stained for Brn3a. Cell count for each microbead-injected eye was normalized against the contralateral BSS-injected eye to determine percent survival of Brn3a+ cells. Representative images are shown on the right. Scale bar = 50μm. (B) Retinal flatmounts were stained for Rbpms. Cell count for each microbead-injected eye was normalized against the contralateral BSS-injected eye to determine percent survival of Rbpms+ cells. Representative images are shown on the right. Scale bar = 50μm. n=13 NSS, n=15 NLY01. All data presented as mean ± SEM. Mann-Whitney U test, ***p<0.001, ****p<0.0001 vs. BSS WT.

## DISCUSSION

Glaucoma is one of the most common neurodegenerative diseases with potentially severe visual implications. Therapies to slow disease progression are currently limited to IOP reduction through both medical and surgical means. Unfortunately, successful reduction of IOP does not prevent disease progression in a large number of patients. New therapies targeting other risk factors for glaucoma are needed to prevent irreversible vision loss. We examined the role of A1 reactive astrocytes in the microbead-induced ocular hypertension mouse model of glaucoma. IL-1α, TNF-α, and C1q have been shown to be necessary and sufficient to drive the formation of A1 astrocytes. Following induction of ocular hypertension, CD11b+ CD11c+ cells, a population enriched for infiltrating macrophages in addition to some resident retinal microglia, initially contributed to IL-1α, TNF-α, and C1q production. The contribution of resident microglia (CD11b+ CD11c−) to IL-1α, TNF-α, and C1q production was insignificant until days to weeks after ocular injection. Together, these three cytokines triggered the formation of A1 astrocytes as demonstrated by upregulation of A1-specific transcripts and C3 production in an ACSA2+ retinal cell population (astrocytes and Müller cells). Treatment with the GLP-1R agonist, NLY01, reduced microglial and macrophage production of IL-1α, TNF-α, and C1q, decreased A1 astrocyte conversion, and protected against RGC death in this mouse model of glaucoma.

Recently published transcriptomic data from the DBA/2J mouse model of glaucoma suggested that early inflammation is driven by infiltrating macrophages (CD11b+ CD11c+), while resident microglia (CD11b+ CD11c) initially adopt an anti-inflammatory gene expression pattern (Tribble et al., 2019). Our results support this conclusion, by demonstrating IL-1α, TNF-α, and C1q upregulation by CD11b+ CD11c+ cells before CD11b+ CD11c− cells. Retinal macrophage infiltration has previously been implicated in glaucoma. Macrophages have been observed in sections of human glaucomatous retina and optic nerve in both mild and severe cases (Margeta et al., 2018). Previous work has also shown that progression of visual field loss in normotensive glaucoma is associated with increased systemic levels of macrophage chemoattractant protein-1 (MCP-1) (Lee et al., 2017). Although the presence of the blood-retina barrier confers a degree of immune-privilege to the retina and likely limits the effects of circulating macrophages, disruption of the blood-retina barrier has been observed in diseases of ocular inflammation (Daruich et al., 2018; Kaur et al., 2008; Kokona et al., 2018; Vecino et al., 2016). A significant fraction of glaucomatous retinas exhibit focal bleeds in the nerve fiber layer surrounding the optic nerve head. These so-called Drance hemorrhages represent disruption of the blood-retina barrier, and present an opportunity for blood-borne immune cells, such as macrophages, to enter the retina (Williams et al., 2017). Together these data provide an impetus for additional work characterizing the role of infiltrating macrophages and the integrity of the blood-retina barrier in glaucoma.

Following microbead injections, the time course of IL-1α, TNF-α, and C1q upregulation closely corresponded with the trajectory of IOP increase, lending support to elevated IOP as the initiator of inflammation. Despite a return to normal IOP, pro-inflammatory cytokines remained upregulated at 6 weeks post-injection, suggesting that the inflammatory pathway remained active beyond the inciting elevation of IOP. In human glaucomatous eyes, a reduction in IOP, whether by pharmacological or surgical means, is not always sufficient to prevent further RGC degeneration. Our data suggest that persistent inflammation after normalization of IOP may well contribute to these refractory cases, presenting an additional therapeutic opportunity for patients who have exhausted therapies rooted in IOP reduction.

A1 astrocytes upregulate complement component 3 (C3) (Liddelow et al., 2017), and C3 inhibitors were shown to reduce RGC cell death in the DBA/2J mouse model of glaucoma (Bosco et al., 2018). We demonstrated elevated C3 production following A1 activation and decreased C3 production following NLY01 inhibition in our glaucoma model. Importantly, C3 is not the only source of toxicity from A1 astrocytes (Liddelow et al., 2017). While C3 inhibition may confer some RGC protection, prevention of A1 astrocyte formation may confer more complete protection against eIOP.

Within two weeks of microbead injection, we showed that pro-inflammatory infiltrating macrophages released IL-1α, TNF-α and C1q to trigger A1 astrocyte transformation. Blocking this pathway using NLY01 rescued RGCs from eIOP-induced death measured at 6 weeks post-injection. Previously, it has been demonstrated that RGC loss does not occur in this model until after 4-6 weeks of prolonged eIOP (Calkins et al., 2018; Ito et al., 2016; Sappington et al., 2009). This raises the question of whether RGC-autonomous mechanisms of cell stress must act in concert with the non-cell autonomous mechanism of retinal inflammation to trigger RGC death. Loss of one of these two pathways, conferred by NLY01 administration in our model, was sufficient to rescue RGCs. Consistent with this, neuronal injury, either via optic nerve crush or eIOP, has been shown to be a necessary precursor for astrocyte-mediated neuroinflammatory injury (personal communication, Shane Liddelow). Targeting RGC-autonomous mechanisms of stress, such as by lowering IOP, in combination with retinal inflammation may provide a synergistic effect to rescue additional cells.

NLY01 reduced microglial and macrophage activation and prevented A1 astrocyte formation. NLY01 belongs to a family of similar glucagon-like peptide 1 receptor (GLP-1R) agonists. NLY01 is a long-acting GLP-1R agonist that efficiently penetrates the blood-brain barrier (Yun et al., 2018). In mouse models of Parkinson’s disease, NLY01 concentrations in the brain were higher than in WT animals. The authors attributed this increase in NLY01 concentration to blood-brain barrier breakdown present in the Parkinson’s disease mouse models (Yun et al., 2018). Similarly, blood-retina barrier disruptions in glaucoma (Williams et al., 2017) may serve a therapeutic benefit by providing a gateway for systemically administered therapeutics to access the retina and the optic nerve. This further highlights the need to characterize the state of the blood-retina barrier in glaucoma.

Previous work has shown that NLY01’s effects on CD11b+ cells are not limited to reductions in levels of IL-1α, TNF-α, and C1q. Rather, NLY01 treatment reduced microglia density and relative IBA1 levels in a mouse model of Parkinson’s disease, suggesting a broader immunosuppressive effect (Yun et al., 2018). It is therefore possible that the reduction in RGC death we observed following NLY01 treatment can be partially attributed to a reduction in microglial and macrophage reactivity that is independent of the drug’s effect on astrocyte phenotype. This is consistent with previous data from other groups demonstrating that reduction in CD11b+ cell reactivity protects RGCs from the effects of eIOP (Williams et al., 2017).

The safety of NLY01 in humans is currently being tested in ongoing clinical trials for Parkinson’s disease. NLY01 belongs to a class of therapeutics, GLP-1R agonists, that have been used in the clinic for more than 15 years. During that time, GLP-1R agonists have demonstrated a favorable safety profile in the long-term treatment of type 2 diabetes mellitus (Aroda, 2018). Diabetes is a known risk factor in glaucoma (Khan et al., 2016), and GLP-1R’s wide usage in diabetes treatment presents an opportunity to retrospectively evaluate its effects among patients with glaucoma and type 2 diabetes in a large scale observational study. The findings of that study could provide evidence as to whether existing GLP-1R agonists reduce glaucoma incidence or progression among patients with diabetes.

Glaucoma is a group of diseases with disparate but often interlinking etiologies. Our data highlight neuroinflammation as a mechanism of glaucomatous damage and demonstrated rescue by NLY01 through the drug’s ability to decrease retinal inflammation. Current therapies targeting IOP are not sufficient to prevent visual loss in many glaucoma patients. NLY01, or more broadly the class of GLP-1R agonists, may well be fruitful avenues for future exploration.

## MATERIALS & METHODS

### Experimental animals

All mice were adult (>3 months old), age-, strain- and sex-matched. C57BL6/J (WT) mice were obtained from Jackson labs (Stock Number 000664, Bar Harbor, ME). *Il1a−/−; Tnf−/−*, *C1qa−/−*, and *Il1a−/−; Tnf−/−; C1qa−/−* (TKO) animals were generously donated by Ben Barres (Stanford University). All animals were fed *ad libitum* and maintained on a 12 h/12 h light/dark cycle in a University of Pennsylvania vivarium. All procedures were approved by the Institutional Animal Care and Use Committee of the University of Pennsylvania and complied with the ARVO Statement for the Use of Animals in Ophthalmic and Vision Research. NLY01 was obtained through a material transfer agreement with Neuraly (Baltimore, MD). A cohort of mice was treated with either twice weekly subcutaneous injections of NLY01 (5 mg kg^−1^ per injection) or with an equivalent amount of normal saline solution (L21819; Fisher Scientific, Waltham, MA).

### Anterior Chamber Injection and Intraocular Pressure (IOP) Measurement

The microbead occlusion model was used to induce elevated intraocular pressure as described previously (Cui et al., 2020). Briefly, mice were anesthetized with intraperitoneal injections of ketamine (80 mg/kg, Par Pharmaceutical, Spring Valley, NY), xylazine (10 mg/kg, Lloyd Inc., Shenandoah, IA), and acepromazine (2 mg/kg, Boehringer Ingelheim Vetmedica, Inc. St. Joseph, MO). Pupils were dilated with topical 1% tropicamide and 2.5% phenylephrine (Akorn, Inc., Lake Forest, IL). Proparacaine anesthetic eye drop at a concentration of 0.5% (Sandoz, Inc., Princeton, NJ) were applied immediately prior to injection. Injection micropipettes were pulled from glass capillaries to a final diameter of ~ 100 μm and connected to a microsyringe pump. Using a micromanipulator for positioning, one eye of the mouse was injected with 1.5 μl of sterile 4.5 μm-diameter magnetic microbeads (1.6 × 10^6^ beads/μL of balanced salt solution; Thermo Fisher Scientific, Waltham, MA) at a location < 1 mm central to the limbus, while the other eye was injected with an equivalent volume of balanced salt solution (Alcon Laboratories, Inc., Fort Worth, TX). A hand-held magnet was used to target the beads into the drainage angle. After injection, 0.5% moxifloxacin antibiotic drops (Sandoz, Inc., Princeton, NJ) was applied to the eye. IOP was measured between 8 and 11 a.m. using the Icare TONOLAB tonometer (Icare TONOVET, Vantaa, Finland). An average of three measurements/eye was used.

### Retinal cell sorting

ACSA2+, CD11b+ CD11c−, and CD11b+ CD11c+ cells were isolated from adult murine retinas using the Miltenyi Adult Brain Dissociation Kit (Miltenyi Biotec, Bergisch Gladbach, Germany, Miltenyi 130-107-677) and MACS magnetic cell separation system. Briefly, mice were anesthetized, euthanized, and whole neurosensory retinas were harvested. Retinal tissue was dissociated in manufacturer provided enzyme mixtures using the gentleMACS dissociator pre-set program: 37C_ABDK_02. Retinal cell suspensions were passed through a 70μm filter (130-098-462, Miltenyi Biotech) and resuspended in debris removal solution. Following debris removal, retinal cells were resuspended and incubated in red blood cell removal solution for 10 mins at 4°C. The following isolation steps were performed: (1) positive selection for ACSA2 (astrocyte and Muller cell enriched, 130-097-678, Miltenyi Biotech), (2) the remaining negative selection pool was then subjected to positive selection for CD11b (microglia and macrophage enriched, 130-093-636 Miltenyi Biotech), and (3) the CD11b+ cells were then selected against CD11c (130-125-835 Miltenyi Biotech). This resulted in three cell populations: (1) ACSA2+, (2) ACSA2− CD11b+ CD11c−, and (3) ACSA2− CD11b+ CD11c+. Positive and negative selection steps used MACS magnetic separation LS columns (130-042-401, Miltenyi Biotech) according to the manufacturer’s protocol. Cells were subsequently used for RNA isolation using the RNeasy Mini Kit (Qiagen, Valencia, CA), according to the manufacturer’s protocol.

### Quantitative PCR (qPCR)

RNA isolation was performed according to the manufacturer’s protocol (RNeasy kit; Qiagen). cDNA was synthesized with reverse transcription agents (TaqMan Reverse Transcription Reagents, Applied Biosystems) according to the manufacturer’s protocol. Realtime qPCR (TaqMan; ABI, Foster City, CA) was performed on a sequence detection system (Prism Model 7500; ABI) using the ΔΔCT method, which provided normalized expression values (normalized against *Gapdh*). All reactions were performed in technical triplicates (three qPCR replicates per qPCR probe).

### Enzyme-linked immunosorbent assays

ELISA kits were used to measure protein levels of IL-1α (BMS627, Thermofisher Scientific), TNF-α (BMS607-3, Thermofisher Scientific), C1q (LS-F55223-1, LifeSpan BioSciences), and C3 (ab157711, Abcam) according to the manufacturer’s protocol. Briefly, mouse retinas were collected and homogenized in phosphate-buffered saline containing the protease inhibitor phenylmethylsulfonylfluoride (100 lM; EMD, Gibbstown, NJ, USA). Assays were performed according to the manufacturers’ protocol. Protein levels were determined by comparing the absorbance produced by the samples with that of a calibration curve.

### Preparation of retinal flatmounts, immunofluorescence & cell counting

RGC quantification and immunolabeling of flat-mounted retinas were performed as previously described (Cui et al., 2020). Briefly, eyes were enucleated and fixed in 4% paraformaldehyde. Retinas were isolated, mounted on glass slides and serially washed with 0.5% Triton X-100 in phosphate-buffered saline (PBS). Flat-mounted retinas were incubated overnight at 4°C with antibodies against RNA-binding protein with multiple splicing (RBPMS; EMD Millipore Corp., Burlington, MA) diluted 1:500 in blocking buffer (2% bovine serum albumin, 2% Triton X-100 in PBS) and brain-specific homeobox/POU domain protein-3a (Brn3a; Synaptic Systems, Goettingen, Germany) diluted 1:1000 in blocking buffer. The following day, retinas were washed and incubated with Alexa Fluor 488 donkey anti-rabbit IgG (Invitrogen, Carlsband, CA; 1:1000 in blocking buffer) and Cy3 goat anti-guinea pig IgG (Abcam, Cambridge, MA; 1:500 in blocking buffer) secondary antibodies for 3 hours at room temperature. After serial washes, flat-mounts were cover-slipped with Vectashield mounting medium containing DAPI (Vector Laboratories Inc., Burlingame, CA). For each flat-mount, 12 standardized photomicrographs were taken at 1/6, 3/6, and 5/6 distance from the center of the retina at 40x magnification by a masked operator. A masked counter quantified the number of RBPMS- and Brn3a-positive cells in each 40x field (0.069 mm^2^) using Nikon Elements analysis software version 4.1 (Nikon Instruments, Melville, NY, USA). The average number of cells in the 12 standardized photomicrographs from the microbead-injected eye was normalized to the average number of cells in the 12 standardized photomicrographs from the BSS-injected eye to calculate percent survival in the microbead injected eye with eIOP.

### Statistical Analysis

All statistical analyses were done using GraphPad Prism 8.0 software. Data was analyzed either by one-way ANOVA followed by Tukey’s multiple comparisons test for comparing between three or more samples, or Mann-Whitney U test for comparing between two samples with 95% confidence without assuming a gaussian distribution. Power calculations were performed using G* Power Software V 3.1.9.7 (Faul et al., 2007). Group sizes were calculated to provide at least 80% power with the following parameters: probability of type I error (0.05), effect size (0.25).

## Acknowledgements

B. Barres donated *Il1a−/−;Tnf−/−* DKO, *C1qa−/−* single knockout, and *Il1a−/−;Tnf−/−;C1qa−/−* TKO mice. P. Williams and M. Margeta provided helpful insight on the role of CD11b+ CD11c+ and CD11b+ CD11c− cells in the retina. JLD is funded by R01EY015240, R01EY028916, Research to Prevent Blindness, The F.M. Kirby Foundation, The Paul and Evanina Bell Mackall Foundation Trust, A gift in memory of Lee F. Mauger, MD. QNC is funded by K08EY029765, K12EY015398, American Glaucoma Society. JS was supported by T32GM007170 during the conceptualization and initiation of this work.

## Author Contributions

Conceptualization, JKS and QNC. Methodology, JKS, SG, and QNC. Formal Analysis, JKS. Investigation, JKS, MA, AB, KU, AGR, and QNC. Writing – Original Draft, JKS and QNC. Writing – Review & Editing, JKS, MA, AB, SG, KU, AGR, JD and QNC. Visualization, JKS. Supervision, JD and QNC. Project Administration and Funding Acquisition, JD and QNC.

## SUPPLEMENTAL FIGURES

**Supplemental Figure 1.**
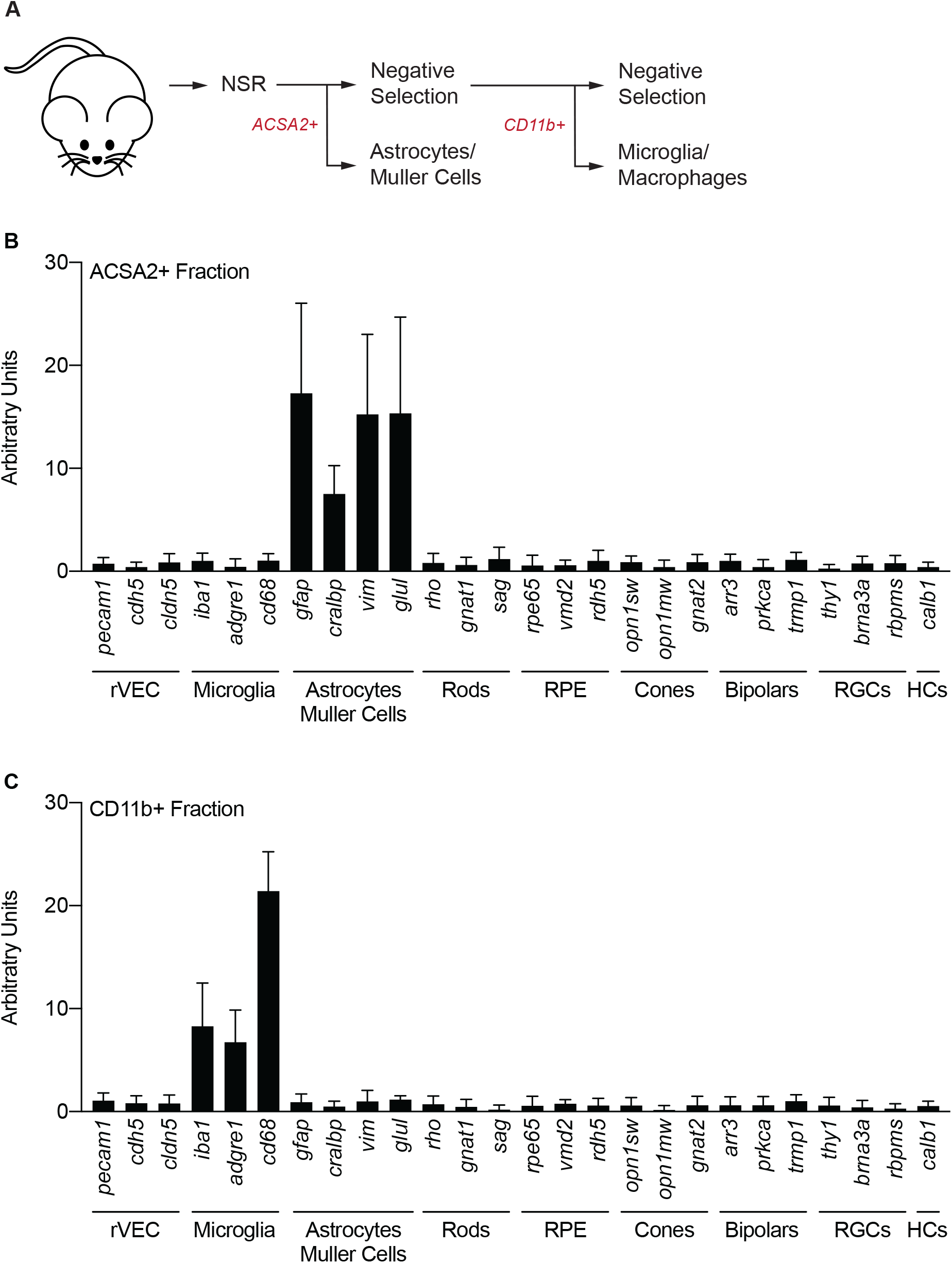
Cell sorting paradigm. C57BL6/J (WT) mice were euthanized and neurosensory retina (“NSR”) was isolated and dissociated for cell sorting. (A) Cell isolation protocol using magnetic cell sorting. (B) qPCR measurements of cell-type specific markers in ACSA2+ cells. (C) qPCR measurements of cell-type specific markers in CD11b+ cells. n = 8 eyes. All data presented as mean ± SEM.

**Supplemental Figure 2.**
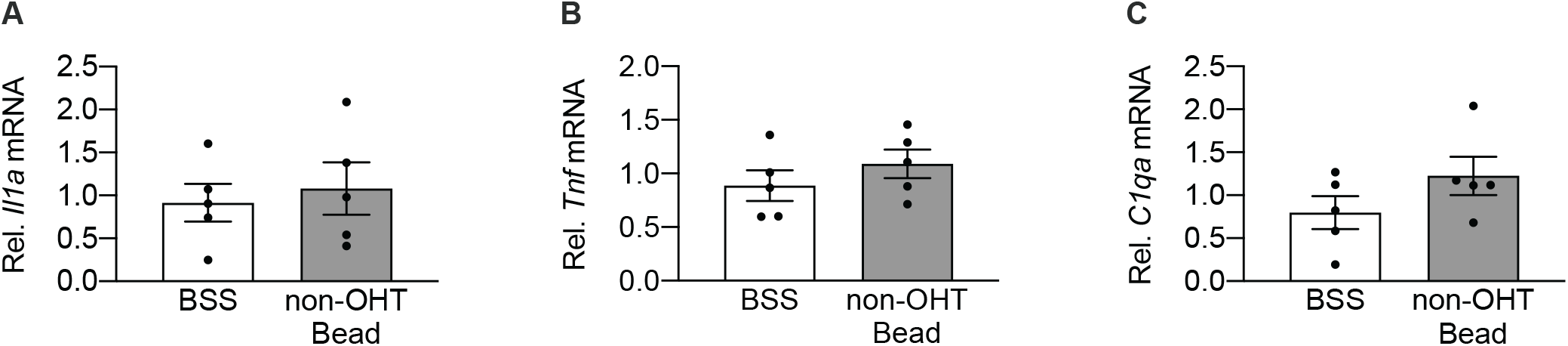
Microbead-injection alone does not induce CD11b+ production of *Il1a*, *Tnf*, or *C1qa*. C57BL6/J (WT) mice were injected either with microbeads (left eye), to increase intraocular pressure (IOP), or with BSS (right eye). IOP was monitored weekly. Bead eyes that did not have an IOP increase of 6 mmHg or greater within 2 weeks of injections were termed “non-OHT Bead” and excluded from eIOP studies. (A-C) CD11b+ cells were isolated from neurosensory retina 42 days after injection. qPCR was performed to measure *Il1a* (A), *Tnf* (B), and *C1qa* (C) mRNA levels. n=5 eyes per condition. All data presented as mean ± SEM.

**Supplemental Figure 3.**
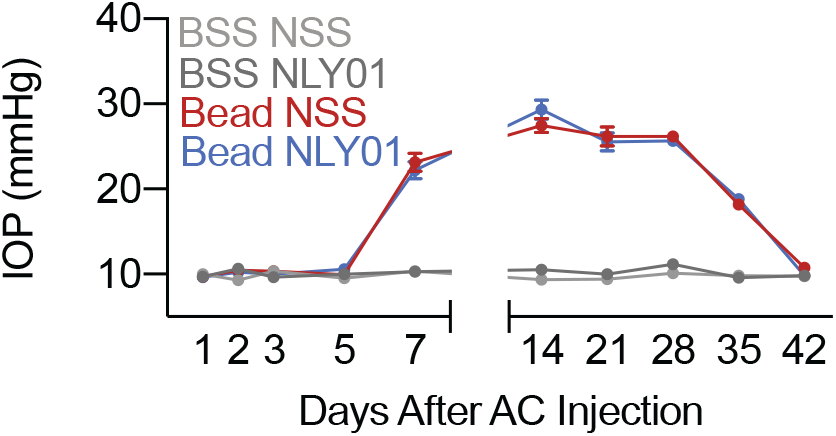
NLY01 does not affect intraocular pressure. C57BL6/J (WT) mice were injected either with microbeads (“Bead”, left eye), to increase intraocular pressure (IOP), or with BSS (right eye). Following intraocular injections, mice were randomized to twice weekly subcutaneous NLY01 (5 mg kg^−1^ per injection) or normal saline solution (NSS). IOP measurements across the duration of the study. (n=25 eyes per condition per treatment)

